# The Envelope Protein of SARS-CoV-2 Inhibits Viral Protein Synthesis and Infectivity of Human Immunodeficiency Virus type 1 (HIV-1)

**DOI:** 10.1101/2022.03.23.485576

**Authors:** Wyatt Henke, Hope Waisner, Sachith Polpitiya Arachchige, Maria Kalamvoki, Edward Stephens

## Abstract

The human coronavirus SARS-CoV-2 encodes for a small 75 amino acid transmembrane protein known as the envelope (E) protein. The E protein forms an ion channel, like the viroporins from human immunodeficiency virus type 1 (HIV-1) (Vpu) and influenza A virus (M2). Here, we analyzed HIV-1 virus infectivity in the presence of four different β-coronavirus E proteins. We observed that the SARS-CoV-2 and SARS-CoV E proteins reduced HIV-1 yields by approximately 100-fold while MERS-CoV or HCoV-OC43 E proteins restricted HIV-1 infectivity to a lesser extent. This was also reflected in the levels of HIV-1 protein synthesis in cells. Mechanistically, we show that that the E protein neither affected reverse transcription nor genome integration. However, SARS-CoV-2 E protein activated the ER-stress pathway associated with the phosphorylation of eIF-2α, which is known to attenuate protein synthesis in cells. Finally, we show that these four E proteins and the SARS-CoV-2 N protein did not significantly down-regulate bone marrow stromal cell antigen 2 (BST-2) while the spike (S) proteins of SARS-CoV and SARS-CoV-2, and HIV-1 Vpu efficiently down-regulated cell surface BST-2 expression. The results of this study show for the first time that viroporins from a heterologous virus can suppress HIV-1 infection.

**IMPORTANCE:** The E protein of coronaviruses is a viroporin that is required for efficient release of infectious virus and for viral pathogenicity. We determined if the E protein from four β-coronaviruses could restrict virus particle infectivity of HIV-1 infection. Our results indicate that the E proteins from SARS-CoV-2 and SARS-CoV potently restricted HIV-1 while those from MERS-CoV and HCoV-OC43 were less restrictive. Substitution of the highly conserved proline in the cytoplasmic domain of SARS-CoV-2 E abrogated the restriction on HIV-1 infection. Mechanistically, the SARS-CoV-2 E protein did not interfere with viral integration or RNA synthesis but rather reduced viral protein synthesis. We show that the E protein-initiated ER stress causing phosphorylation of eIF-2α, which is known to attenuate protein synthesis. Companion studies suggest that the E protein also triggers autophagy. These results show for the first time that a viroporin from a coronavirus can restrict infection of another virus.

## INTRODUCTION

The recent introduction of the highly transmissible coronavirus SARS-CoV-2 into the human population has resulted in the COVID-19 respiratory disease and pandemic (**1–4**). This has highlighted the urgent need to better understand the function of viral proteins in replication and virus pathogenesis. The human pathogenic β-coronaviruses (SARS-CoV-2, SARS-CoV, MERS-CoV, HCoV-OC43) contain four structural proteins (S, M, E, and N) (**5, 6**). The spike (S) protein, the largest protein within virions, binds to angiotensin converting enzyme 2 (ACE-2) receptor, and is the major target for neutralizing antibodies. The membrane (M) protein, the most abundant protein of the virion, is thought to form a scaffold in the ER-Golgi intermediate compartment (ERGIC), the site of virus maturation (**7,8**). The nucleocapsid (N) of SARS-CoV binds to the viral RNA and together with E and M can form viral particles (**9**).

The E protein of the β-coronaviruses is the smallest and least abundant virion protein. This transmembrane protein (75-84 amino acids in length) is predicted to have a short N-terminal domain, a hydrophobic transmembrane domain, and a longer cytoplasmic domain. Recent studies indicate that the E protein spans the membrane a single time with the N-terminal domain facing the lumen of the ER/ERGIC/Golgi (**10–12**). While the expression of SARS-CoV E protein is dispensable for coronavirus replication, its deletion results in reduced virus growth, likely due to inefficient assembly (**13–15**). The E proteins of SARS-CoV-2 and SARS-CoV have 94% amino acid identity, differing at only four amino acid positions, while E proteins of MERS-CoV and HCoV-OC43 are more distantly related to SARS-CoV-2 E protein with approximately 37% and 25% amino acid identity, respectively. Interestingly, the E proteins of SARS-CoV-2 and several bat coronavirus isolates (RATG13, ZC45, and ZXC21) are identical.

The E proteins of coronaviruses have similarities to the Vpu protein of HIV-1 with respect to its size, orientation in the membrane, ion channel activity, and functions to enhance virus release (**16–23**). These similarities prompted us to examine the biological properties of the SARS-CoV-2 E protein in the context of HIV-1 replication. Our results indicate that the SARS-CoV-2 E protein potently restricted HIV-1 infection. This restriction was not due to inhibition of viral integration or synthesis of viral RNA but rather correlates with the ability of the SARS-CoV-2 E protein to activate the phosphorylation of the eIF-2α, which is known to shutdown protein synthesis. These results show for the first time that a viroporin (i.e., E protein) from SARS-CoV-2 that enhances virus infection can restrict a heterologous virus (i.e., HIV-1).

## RESULTS

### The E protein of SARS-CoV-2 is expressed within the RER and Golgi complex and potently restricts the release of infectious HIV-1

We first examined the intracellular expression of the SARS-CoV-2 E protein. COS-7 cells were transfected with the vector expressing the SARS-CoV-2 E protein with a C-terminal HA-tag and immunostained for the presence of the E protein (anti-HA and either ER (anti-calnexin) or Golgi (anti-Golgin 97). The results show that the E protein co-localized with markers for the ER (calnexin) and Golgi (Golgin-97) but was not observed at the cell surface (**Fig. 1**) as was previously observed (**11, 24–28**). We next determined if the SARS-CoV-2 E protein would enhance or restrict the level of HIV-1Δ*vpu* or HIV-1 particle infectivity released from cells. HEK293 cells were co-transfected with the empty pcDNA3.1(+) vector, or pcDNA3.1(+) expressing either the SARS-CoV-2 E protein, HSV-1 gD (positive control for restriction) or gD[ΔTMCT] (a truncated form of gD that is secreted from cells and does not restrict) and pNL4-3 or pNL4-3*vpu* (**29, 30**). As previously reported, gD potently restricted the release of infectious HIV-1 while gD[ΔTMCT] did not restrict HIV-1 (**Fig. 2A**). Our results indicated that co-transfection with vectors expressing SARS-CoV-2 E protein and plasmids with either HIV-1 (strain NL4-3) or HIV-1Δ*vpu* genomes resulted in potent restriction of the release of infectious virus, which was like the gD protein of HSV-1(**Fig. 2A**). Immunoprecipitation of gD and SARS-CoV-2 E proteins (anti-HA for the E protein; anti-gD for HSV-1 gD) from the cell lysates revealed that these proteins were expressed well during the co-transfections (**Fig. 2B**).

**Figure 1.**
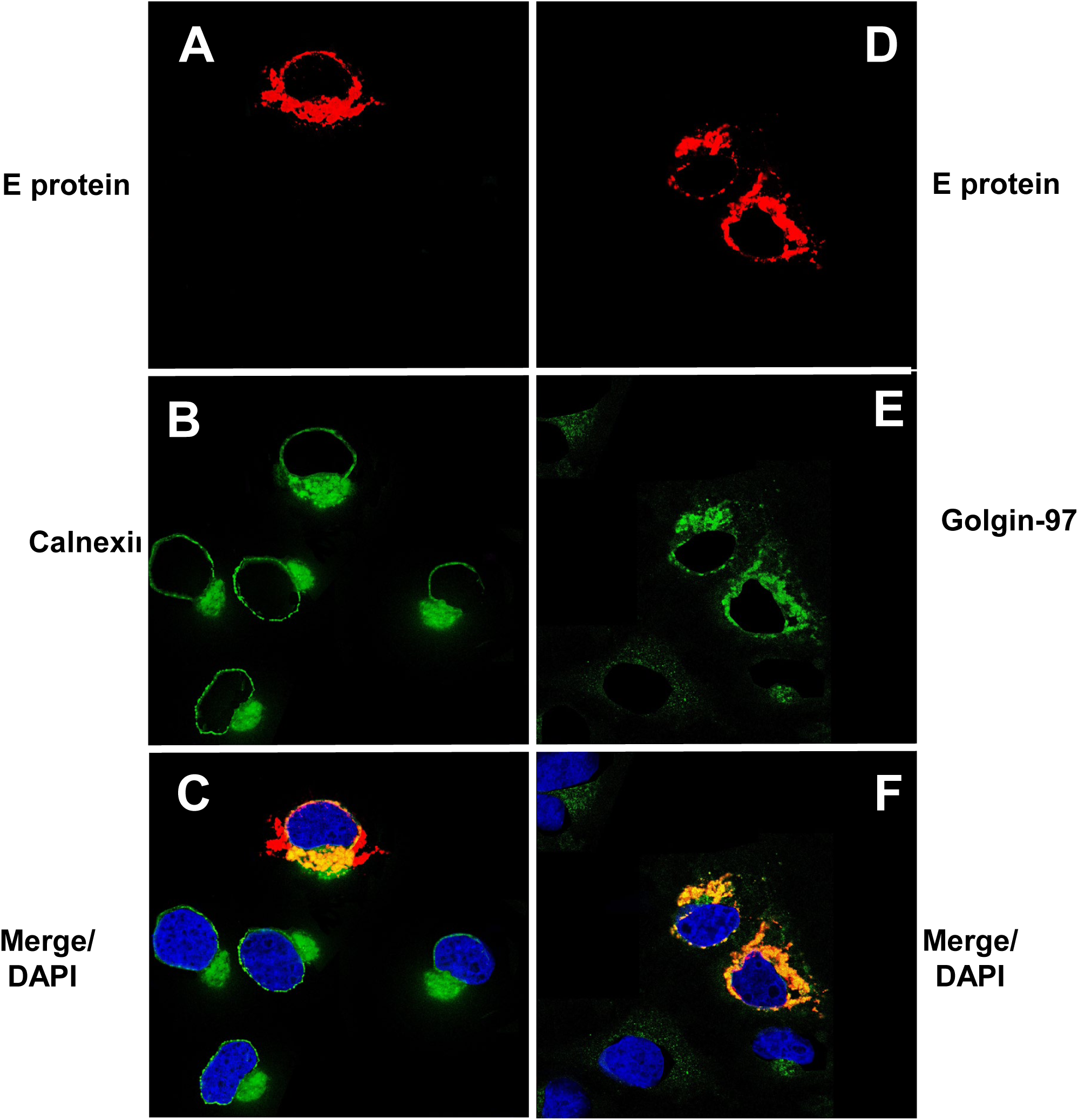
The expression of the SARS-CoV-2 E protein is restricted to intracellular compartments of the cell. COS-7 cells were transfected with the vector expressing the SARS-CoV-2 E-HA protein. At 24 h, cells were fixed, permeabilized and blocked. The cover slips were reacted with a mouse monoclonal antibody against HA-tag and with a rabbit antibody against the ER marker calnexin or trans Golgi network protein Golgin-97 overnight at 4C followed by an appropriate secondary antibody for 1 h. Cells were washed, and counter stained with DAPI (1 μg/ml) for 5 min as described in the Materials and Methods section. The cover slips were mounted and examined using a Leica TCS SPE confocal microscope using a 63x objective with a 2x digital zoom using the Leica Application Suite X LAS X, LASX) software package. A 405nm filter used to visualize the DAPI and a 594nm filter to visualize HA staining and 488 nm filter to visualize the ER or Golgi marker staining. **Panels A-C**. Cells transfected with the vector expressing E-HA and immunostained for the HA tag and calnexin. **Panels D-F**. Cells transfected with the vector expressing E-HA and immunostained for the HA tag and Golgin-97.

**Figure 2.**
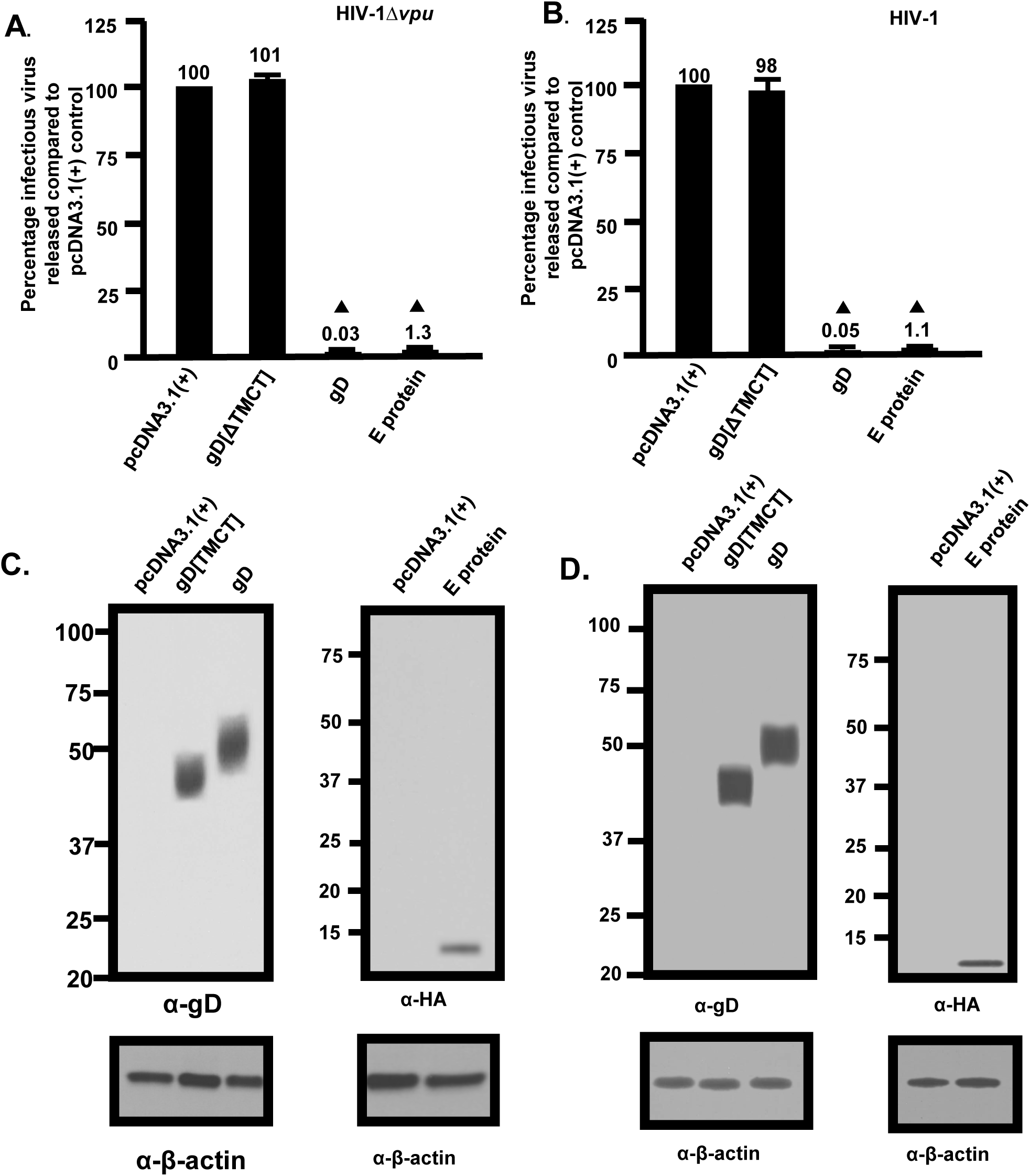
Release of infectious HIV-1Δ*vpu* and HIV-1 in the presence of SARS-CoV-2 E protein. HEK293 cells were co-transfected with either the empty pcDNA3.1(+) vector or vectors expressing the HSV-1 gD[ΔTMCT], HSV-1 gD, or SARS-CoV-2 E protein and pNL4-3 or pNL4-3Δ*vpu*. At 48 h, the culture supernatants were collected, subjected to low-speed centrifugation to remove cellular debris, and levels of infectious virus released into the culture supernatants was determined using TZM-bl cell assays. **Panel A**. The level of HIV-1Δ*vpu* infectivity in the culture medium from cells co-transfected with either the empty pcDNA3.1(+) vector or vectors expressing gD[ΔTMCT], gD, or SARS-CoV-2 E protein and pNL4-3Δ*vpu*. **Panel B**. The level of HIV-1 infectivity in the culture medium from cells co-transfected with either the empty pcDNA3.1(+) vector or vectors expressing gD[ΔTMCT], gD, or SARS-CoV-2 E protein and pNL4-3. **Panel C.** Expression of the gD proteins or E protein from restriction assays in Panel A. **Panel D.** Expression of the gD proteins or E protein from restriction assays in Panel B. Expression of the E protein or gD proteins from restriction assays in Panel A and B normalized with β-actin by immunoprecipitation using an anti-β-actin antibody. All restriction assays in A and B were performed at least four times and statistical differences with the pcDNA3.1(+)/HIV-1 control evaluated using a two-tailed Student’s *t*-test, with *p*<0.01 (▲) considered significant.

### The SARS-CoV-2 E protein did not inhibit HSV-1 infection

We next determined if the restriction of release of infectious HIV-1 was observed in other virus infections. HEK293 cells were transfected with the plasmid expressing SARS-CoV-2 E protein for 24 h followed by infection of cultures with HSV-1 (0.01 pfu/cell). Samples were collected at 3, 24, and 48 h post-infection. The results indicate that SARS-CoV-2 E protein did not interfere with infectious HSV-1 progeny virus production at 24 or 48 h post-infection (**Fig. 3**).

**Figure 3.**
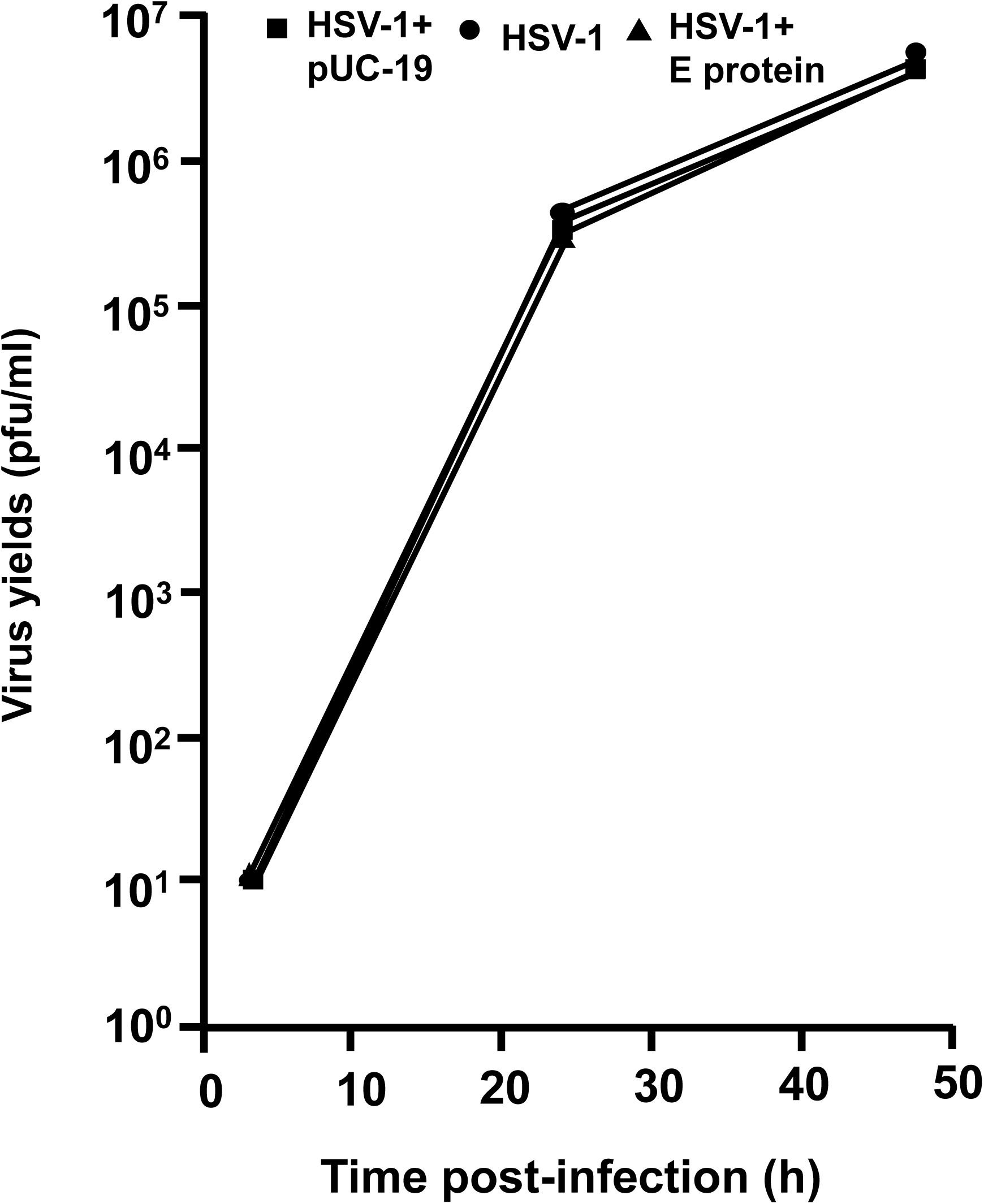
The E protein of SARS-CoV-2 does not restrict the replication of HSV-1. HEK293 cells were left untransfected or transfected with the empty vector (pUC-19) or a vector expressing the SARS-CoV-2 E protein. At 24 h post-transfection, cells were inoculated with HSV-1 (0.01 pfu/cell). Cells were harvested at 24 and 48 h post-infection. The number of plaque forming units were determined using standard plaque assays.

### Restriction of HIV-1 by SARS-CoV, MERS-CoV and HCoV-OC43 E proteins

As the E protein of SARS-CoV-2 appeared to decrease the levels HIV-1 virus released, we constructed pcDNA3.1(+) vectors that expressed the E proteins from three other β-coronaviruses that cause mild to severe pathogenicity in humans (HCoV-OC43, SARS-CoV, MERS-CoV). Like the SARS-CoV-2 E protein, examination of the intracellular expression revealed that these E proteins were primarily localized in the ER and Golgi regions of the cell with no expression at the cell surface (data not shown). We determined if these E proteins could also restrict HIV-1 particle infectivity by transfection of HEK293 cells with vectors expressing the SARS-CoV-2, SARS-CoV, MERS-CoV, or HCoV-OC43 E proteins, HSV-1 gD, or gD[ΔTMCT] and pNL4-3. At 48 h, the culture medium was collected, clarified, and the levels of infectious HIV-1 released determined using TZM.bl cell assays. The presence of gD restricted the release of infectious HIV-1 (0.04%) while the presence of gD [TMCT] did not affect levels of infectious virus released (∼101%) (**Fig. 4A**). The results indicate that the E proteins from SARS-CoV-2 and SARS-CoV potently restricted the release of infectious HIV-1 at 1.3% and 1.4%, respectively. However, MERS-CoV and HCoV-OC43 E proteins were less restrictive at approximately 16% and 37%, respectively, of the pcDNA3.1(+) control (**Fig. 4**). Immunoprecipitation of gD and E proteins from cell lysates from the restriction assays confirmed that the gD and E proteins were expressed (**Fig. 4B**). These results provide additional data on the specificity of the restriction of HIV-1.

**Figure 4.**
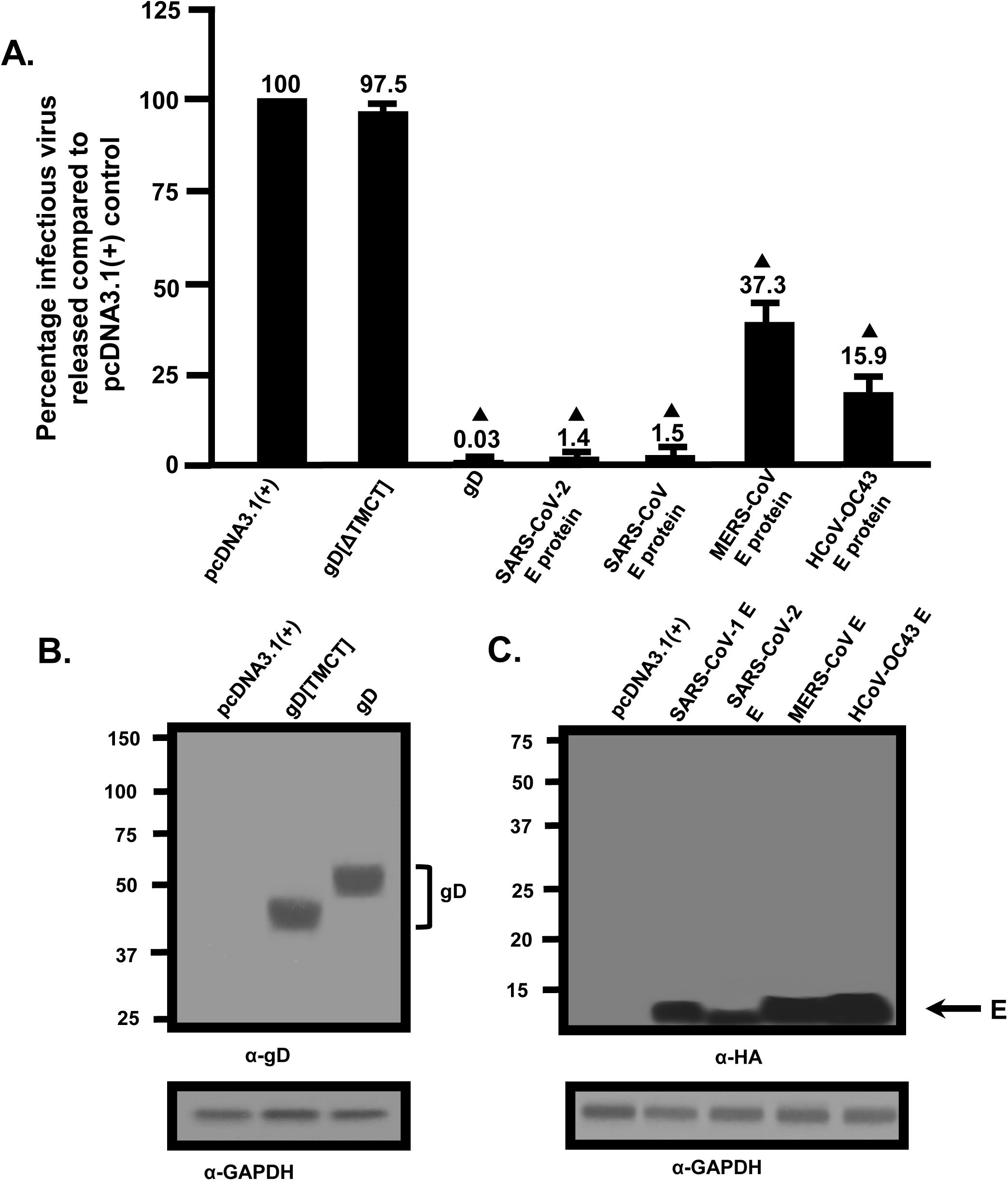
Restriction of the HIV-1 particle infectivity by the SARS-CoV, MERS-CoV, and HCoV-OC43 E proteins. **Panel A**. The infectivity of HIV-1 particles released into the culture medium from cells co-transfected with pcDNA3.1(+) alone, or vectors expressing SARS-CoV-2 E-HA, SARS-CoV E-HA, MERS-CoV E-HA, HCoV-OC43 E-HA, HSV-1 gD, or gD[ΔTMCT] and pNL4-3. **Panel B.** Immunoprecipitation of gD and gD[ΔTMCT] from the cell lysates of the restriction assay in Panel A using an anti-gD antibody. **Panel C.** Immunoprecipitation of the various E proteins from the cell lysates from the restriction assay in Panel A using an anti-HA antibody.

### The SARS-CoV-2 and SARS-CoV E proteins result in decreased biosynthesis of HIV-1 proteins

We examined the mechanism through which the SARS-CoV-2 E protein restricted the release of infectious HIV-1. HEK293 cells were transfected with the empty pcDNA3.1(+) vector or one expressing the SARS-CoV-2 E protein and pNL4-3. At 36 h post-transfection, cells were starved for methionine/cysteine for 2 h and radiolabeled with ^35^S-methionine/cysteine for 16 h. The culture medium was collected and clarified while cell lysates were prepared. HIV-1 and E proteins were immunoprecipitated as described in the Materials and Methods section. Immunoprecipitation of HIV-1 proteins from cell lysates of cells co-transfected with pcDNA3.1(+) and pNL4-3 revealed presence of the gp160 Env, gp120 cleavage product, Gag p55, and p24. Further, gp120 and p24 were immunoprecipitated from the clarified culture medium. In contrast, immunoprecipitation of HIV-1 proteins from lysates prepared from cells co-transfected with the vector expressing SARS-CoV-2 E and pNL4-3 revealed that the Env precursor gp160, the gp120, p55, and p24 were immunoprecipitated at much lower levels as were gp120 and p24 in the clarified culture medium. (**Fig. 5A**). The E protein was detected in the cell lysates but not detected in culture medium (**Fig. 5B**). We also determined if the E proteins from SARS-CoV, MERS-CoV and HCoV-OC43 also affected HIV-1 protein synthesis (**Fig. 5C-H**). The presence of SARS-CoV E also reduced levels of viral proteins in the cell lysates and culture medium compared with the pcDNA3.1(+)/pNL4-3 control (**Fig. 5C-D**) while analysis of MERS-CoV/pNL4-3 and HCoV-OC43/ pNL4-3 co-transfections revealed an intermediate level of viral proteins in the cell lysates and in the culture medium (**Fig. 5E-H**), which correlates with the levels of HIV-1 infectivity.

**Figure 5.**
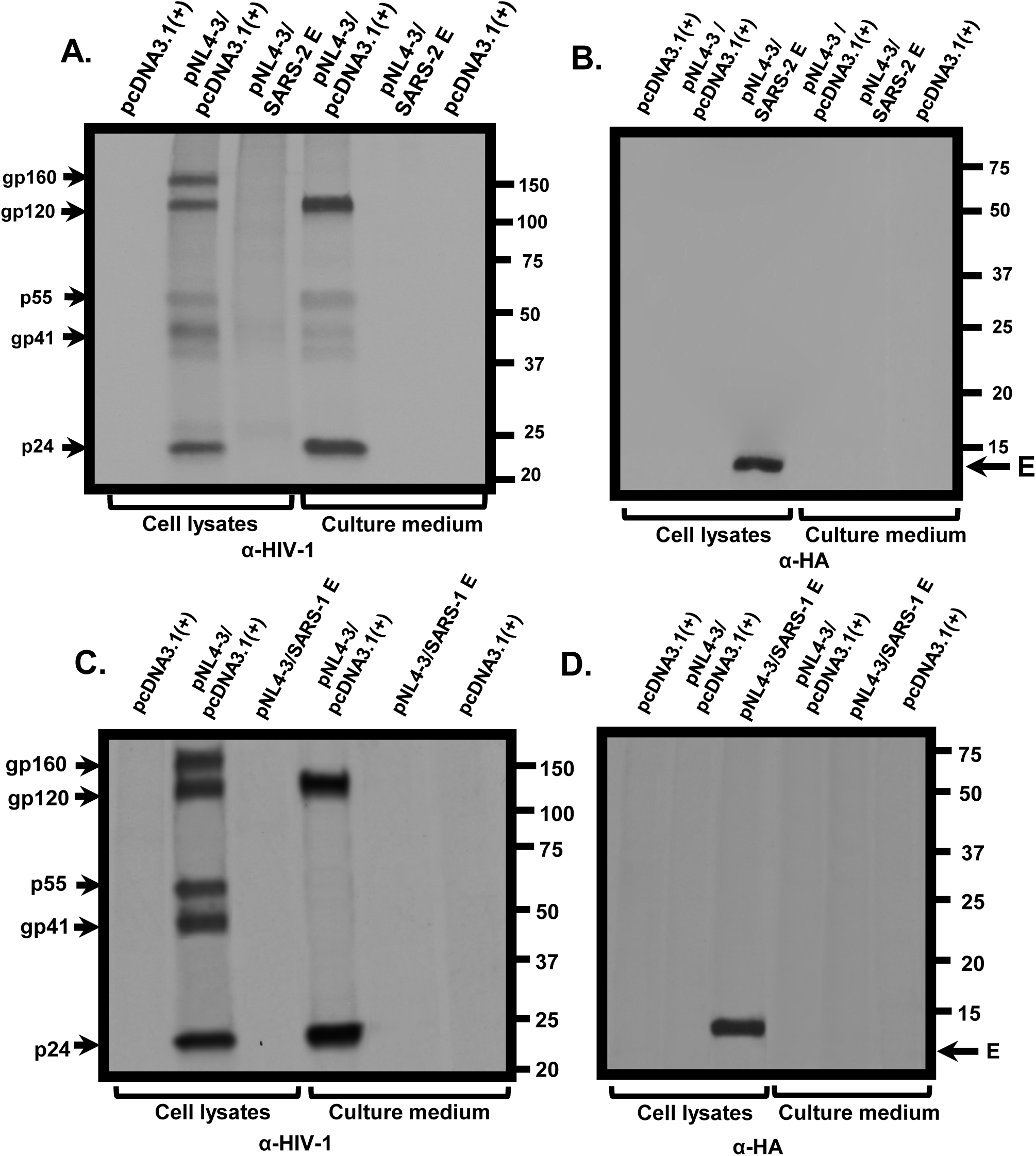
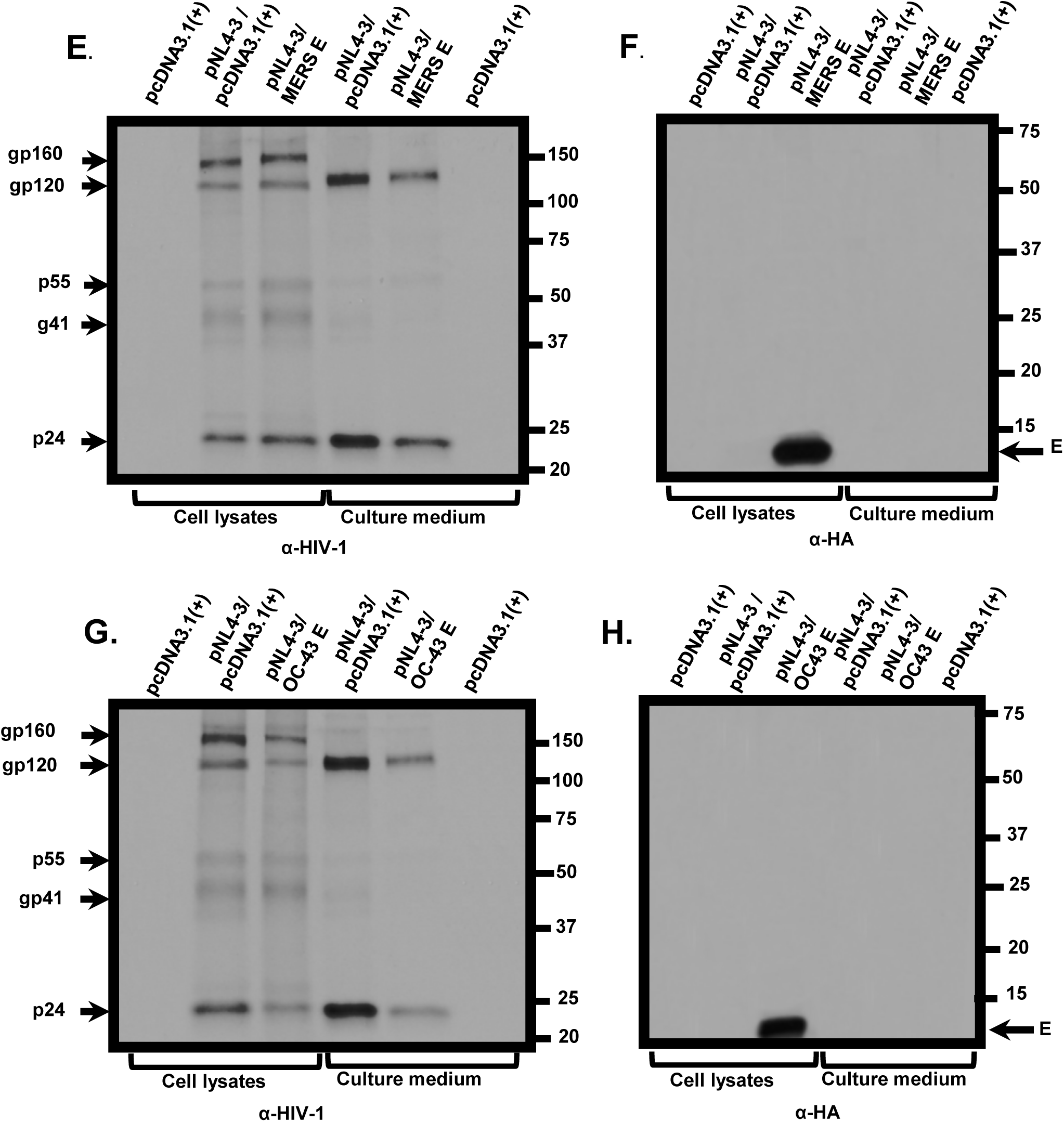
The biosynthesis of HIV-1 proteins is attenuated in the presence of SARS-CoV-2 and SARS-CoV E to higher levels than MERS-CoV and HCoV-OC43 E proteins. HEK293 cells were transfected with either the empty pcDNA3.1(+) or vectors expressing SARS-CoV-2, SARS-CoV, MERS-CoV, or HCoV-OC43 E proteins and pNL4-3. At 30 h post-transfection, cells were starved of methionine/cysteine for 2 h and radiolabeled with ^35^S-methionine/cysteine for 16 h. The culture medium was collected, and cell lysates prepared as described in the Materials and Methods section. HIV-1 proteins were immunoprecipitated using anti-HIV-1 antibodies while the E proteins were immunoprecipitated with antibodies directed against the HA-tag. The immunoprecipitates were collected on protein-A-Sepharose, washed, and boiled in sample reducing buffer. The proteins were separated by SDS-PAGE and visualized using standard radiographic techniques. **Panel A-B.** Immunoprecipitation of HIV-1 proteins from cell lysates (A) and culture medium (B) from HEK293 cells co-transfected with either pcDNA3.1(+) alone, pcDNA3.1(+)/pNL4-3, or pcDNA3.1(+) expressing SARS-CoV-2 E-HA protein/pNL4-3. **Panel C-D.** Immunoprecipitation of E proteins from cell lysates (C) and culture medium (D) from HEK293 cells co-transfected with pcDNA3.1(+) alone, or pcDNA3.1(+)/pNL4-3, or pcDNA3.1(+) expressing the SARS-CoV E-HA/pNL4-3. **Panel E-F.** Immunoprecipitation of HIV-1 proteins from cell lysates and culture medium from HEK293 cells co-transfected pcDNA3.1(+) alone, pcDNA3.1(+)/pNL4-3, or pcDNA3.1(+) expressing the MERS-CoV E-HA/pNL4-3. **Panel G-H.** Immunoprecipitation of HIV-1 proteins from cell lysates and culture medium from HEK2933 cells co-transfected pcDNA3.1(+) alone, pcDNA3.1(+)/pNL4-3, or pcDNA3.1(+) expressing the HCoV-OC43 E-HA/pNL4-3.

### The E protein does not prevent integration of the HIV-1 genome or viral transcription

As the immunoprecipitation assays indicated that HIV-1 protein synthesis was decreased in the presence of the E protein, it suggested that the presence of E protein may have affected an early step in HIV-1 replication. We used Alu-*gag* real time PCR assays to quantify integration of the HIV-1 genome in the presence and absence of the E protein. HEK293 cells were transfected with either the empty pcDNA3.1(+) vector or the vector expressing E-HA. At 24 h, equal infectious units of HIV-1 Env/VSV-G virus were used to infect the transfected HEK293 cells for 48h. The DNA was extracted, and the levels of Alu-gag products determined using the procedures in the Material and Methods section. Our results indicate that integration did not appear to be affected in the presence of the E protein while treatment of cells with raltegravir was effective in preventing integration (**Figure 6A**). We also quantified *tat* gene transcription by real time RT-PCR and the results show that the number of transcripts detected during infection were similar in cells transfected with the empty pcDNA3.1(+) vector or pcDNA3.1(+) expressing E-HA (**Figure 6B**).

**Figure 6.**
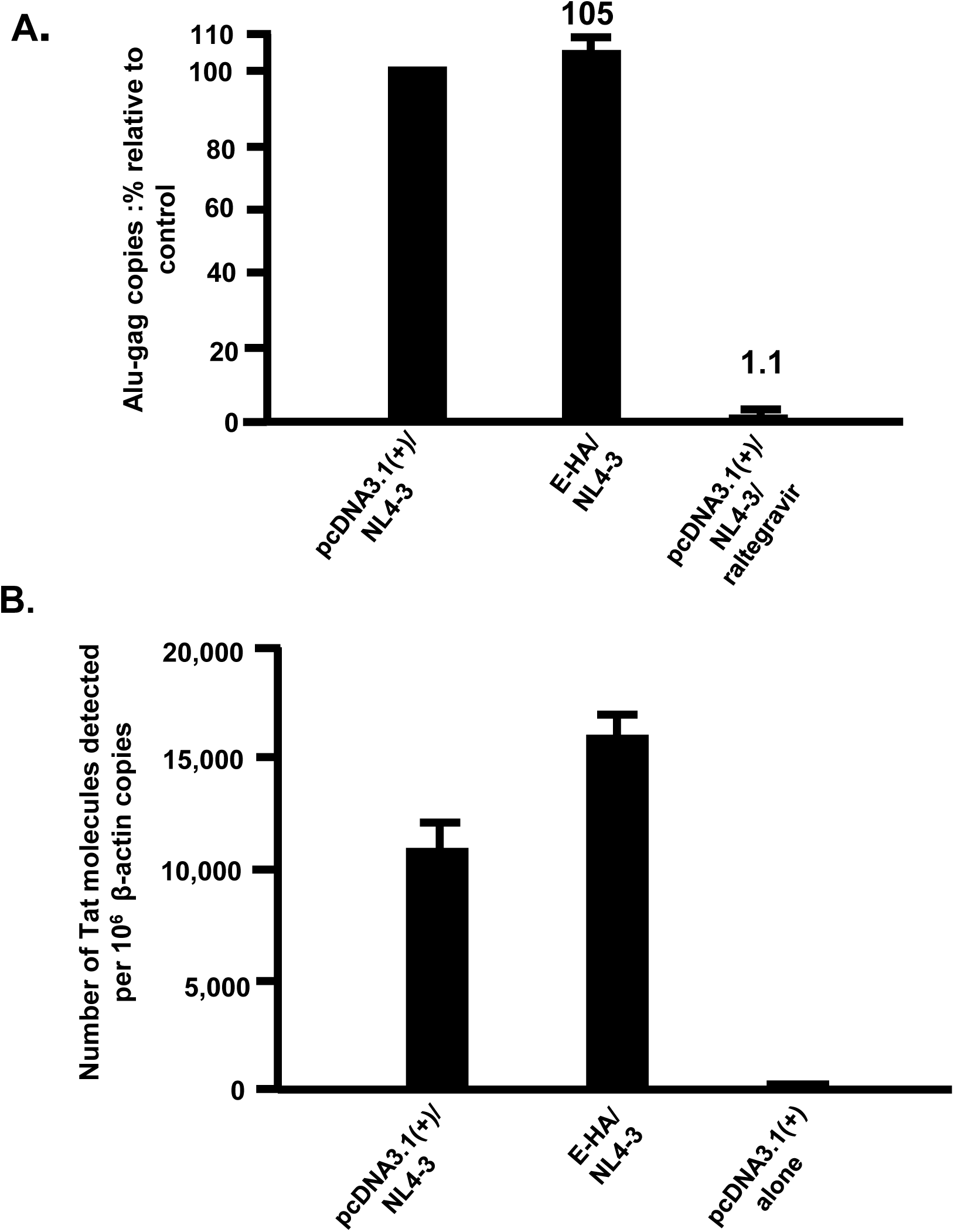
The SARS-CoV-2 E protein did not alter integration or *tat* transcription. **Panel A.** HEK293 cells were transfected with either empty pcDNA3.1(+) vector or one expressing the SARS-CoV-2 E protein. At 24 h, cells were inoculated with equal numbers of infectious units with pseudotyped NL4-3ΔEnv-EGFP/VSV virus as described in the Materials and Methods section. Cells were harvested at 48h post-inoculation, total DNA extracted, and Alu-*gag* products amplified using real time DNA PCR. An additional negative control included treatment of transfected/infected cells with 20μM raltegravir for 2 h prior to inoculation of cultures with pseudotyped virus. **Panel B.** Cells were transfected and infected as in panel A. At 48 h post-transfection, cells were washed, pelleted, and the RNA was extracted. RNA was used in real time RT-PCR reactions to quantify and normalize *tat* RNA levels to β-actin levels.

### Expression of SARS-CoV-2 E protein results in the phosphorylation of eIF-2α

As expression of the SARS-CoV-2 E protein significantly reduced HIV-1 protein expression, we determined if the E protein activated pathways associated with an unfolded protein response due to ER stress such as phosphorylation of eIF-2α, which can inhibit protein synthesis. HEK293 cells were transfected with the vector expressing SARS-CoV-2 E-HA. At 48 h post-transfection (pt) cells were lysed and phosphorylated eIF-2α was analyzed by immunoblots using an antibody against phospho-eIF-2α. In cells transfected with the empty pUC19 vector, little phosphorylated eIF-2α was detected (**Fig. 7**). In contrast, in cells transfected with the vector expressing E-HA phosphorylated eIF-2α was readily detected (**Fig. 7**). These results indicate that expression of E-HA resulted in increased eIF-2α phosphorylation.

**Figure 7.**
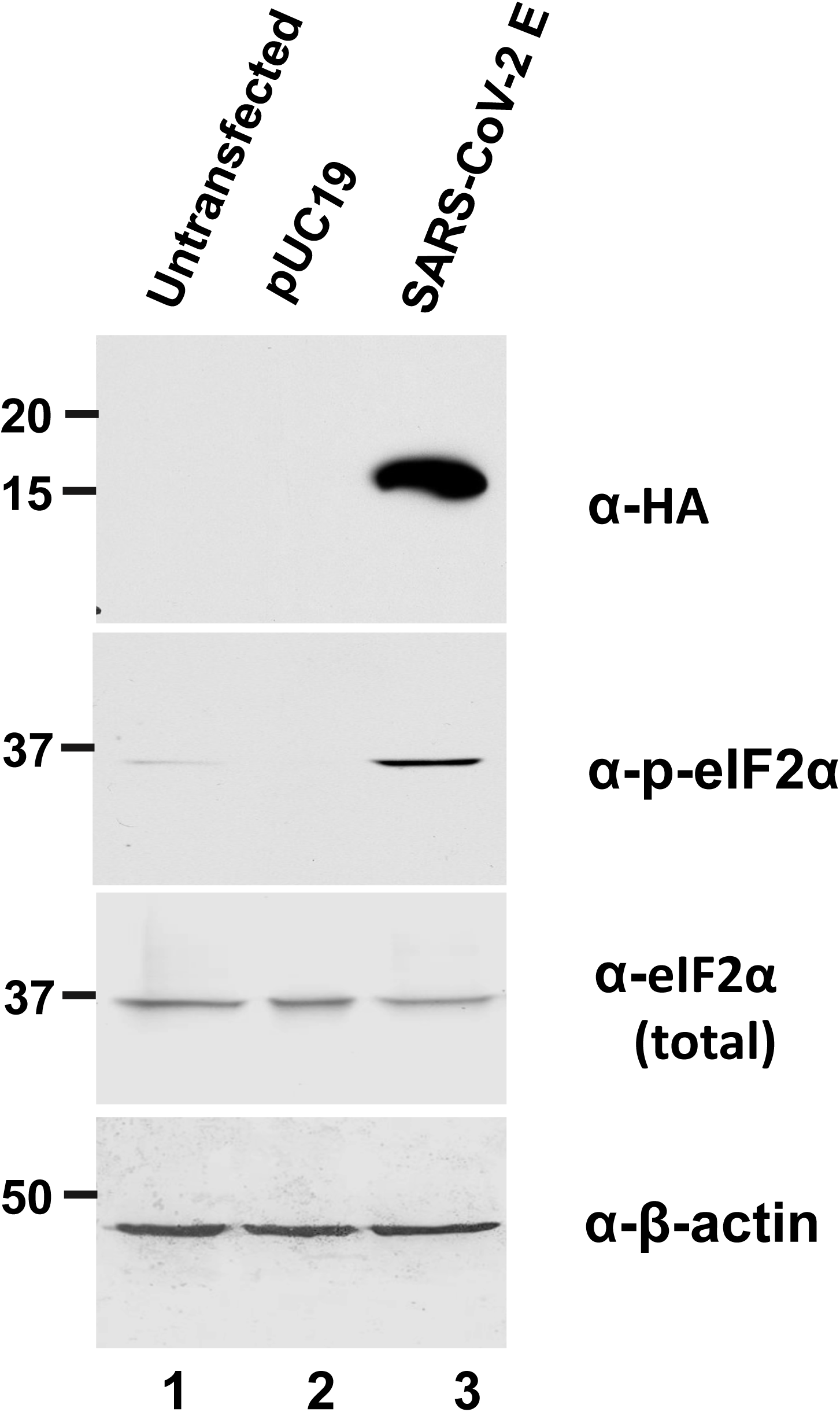
The E protein causes ER-stress and phosphorylation of the eIF-2α. HEK293 cells were either left untransfected or transfected with the empty pUC-19 or pcDNA3.1 (+) expressing the SARS-CoV-2 HA-E proteins. At 48 h post-transfection, cells were lysed, and western blots were used to examine the expression of the E protein (α-HA), phosphorylated eIF2α, total eIF2α, and β-actin (as loading control). Vectors used to transfect cells are at the top of each lane.

### BST-2 down-regulation by E proteins, SARS-CoV-2 S and N proteins

As the HIV-1 Vpu and the coronavirus E proteins have a similar overall structure and are both viroporins, we examined if the E proteins from different coronaviruses, like Vpu, could down-regulate bone marrow stromal antigen 2 (BST-2). Vectors expressing each of the four E proteins, SARS-CoV and SARS-CoV-2 S proteins, SARS-CoV-2 N protein, and the HIV-1 Vpu protein were co-transfected into HEK293 cells with a vector expressing human BST-2. At 48 h post-transfection, cells were immunostained with a monoclonal antibody against BST-2 tagged with Alexa 488 and BST-2 surface expression analyzed by flow cytometry. The mean and median fluorescent intensities were calculated using the FlowJo software program. Our results showed that neither the E proteins nor SARS-CoV-2 N down-regulated BST-2 cell surface expression (**Fig. 8A-B**). However, the SARS-CoV and SARS-CoV-2 S proteins and the HIV-1 Vpu protein significantly down-modulated BST-2 cell surface expression (**Fig. 8A-B**). Analysis of aliquots of cells from the same co-transfections revealed E proteins, S proteins, N and Vpu were all expressed well in co-transfected cells (**Fig. 9C**).

**Figure 8.**
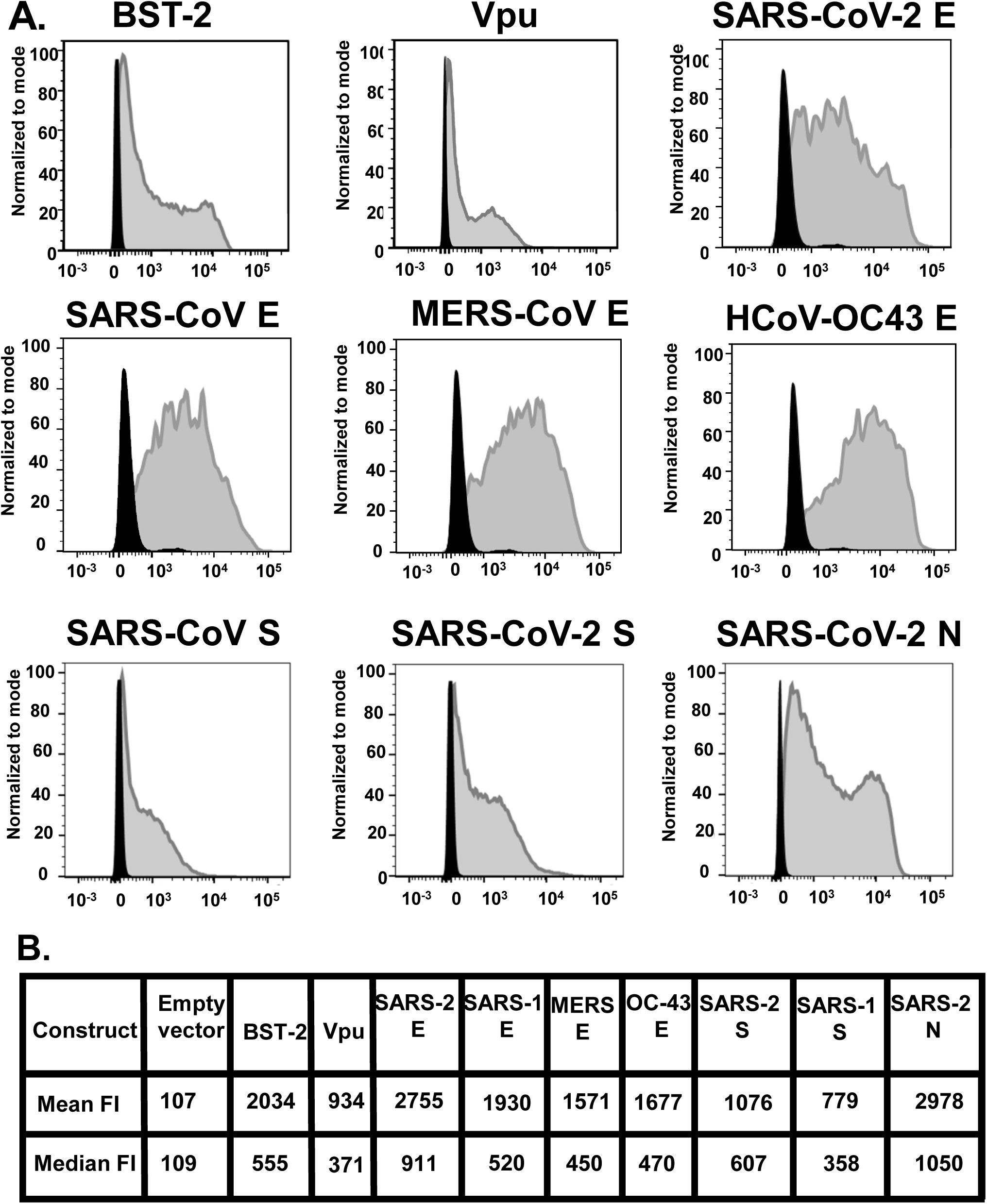
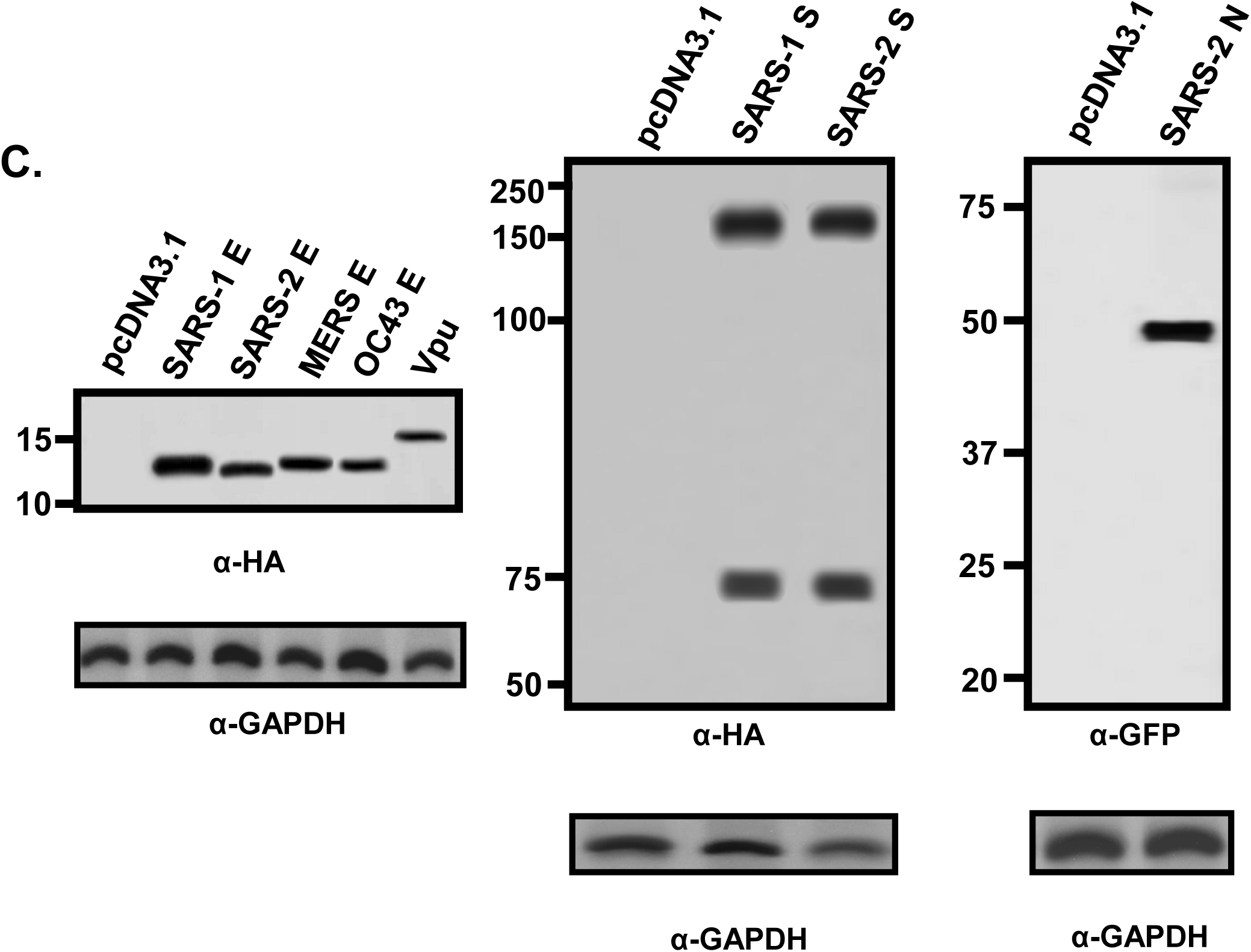
BST-2 down-regulation by E, S and N proteins. HEK293 cells were co-transfected with either the empty pcDNA3.1(+) vector or pcDNA3.1(+) vector expressing each of the four E proteins, SARS-CoV-2 N protein, SARS CoV-2 S protein, and SARS CoV S protein and a vector expressing human BST-2 protein. **Panel A.** At 48 h, the cells were removed by treatment with EDTA/EGTA, stained with a mouse monoclonal antibody directed against BST-2, fixed and subjected to flow cytometric analysis using a BD LSR II flow cytometer. The fluorescent intensities are on the x-axis. **Panel B**. Median and mean fluorescent intensities of the BST-2 on the cells from Panel A. **Panel C.** Aliquots of cells from the same co-transfection were also analyzed for protein expression by immunoblots using antibodies directed against the HA-tag (E proteins, SARS-CoV-2 S protein, and Vpu), an antibody against SARS-CoV S protein, or GFP to monitor SARS-CoV-2 N-GFP expression.

**Figure 9.**
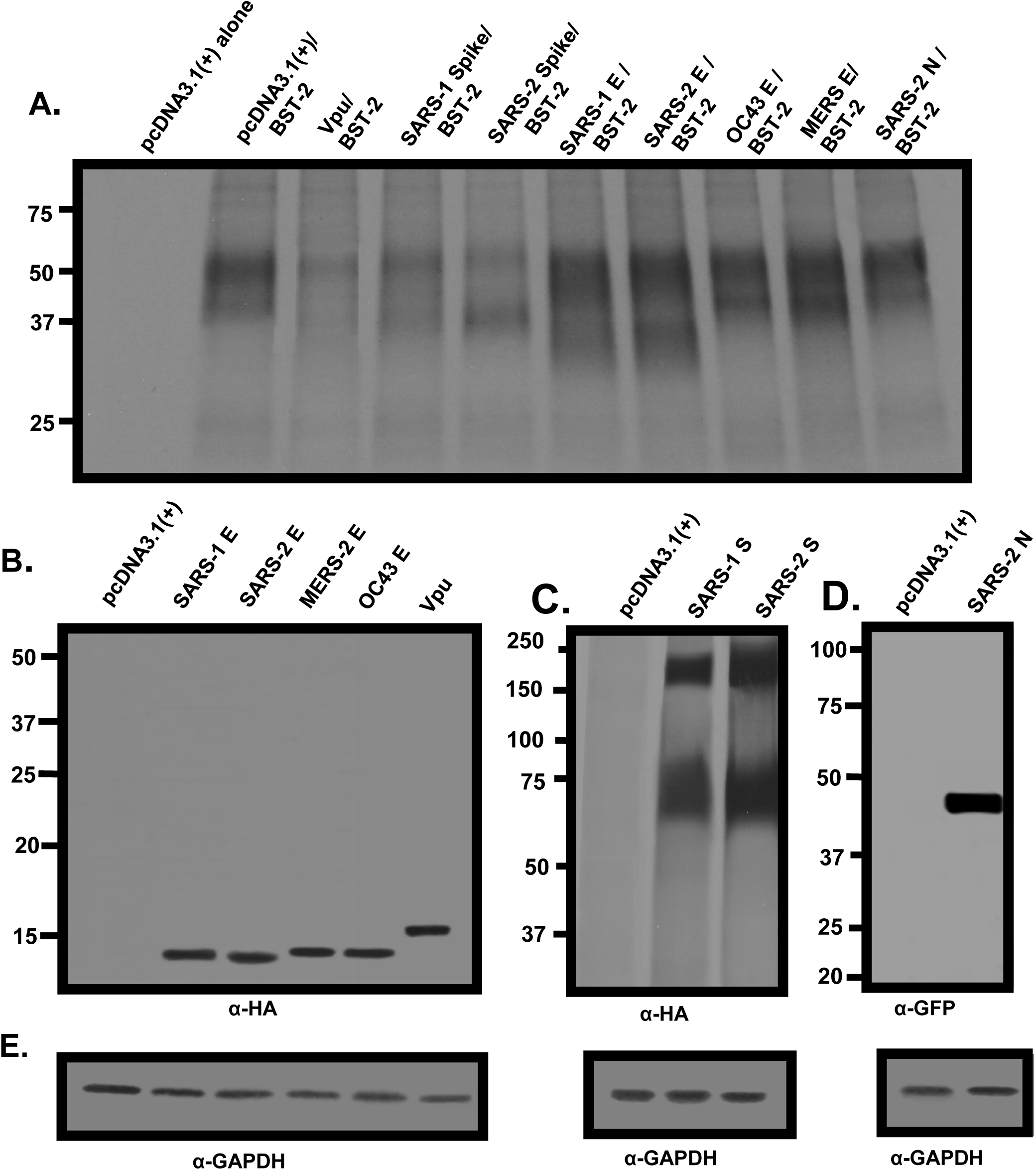
The levels of BST-2 in cells in the presence of coronavirus proteins. HEK293 cells were co-transfected with either the empty pcDNA3.1(+) vector or pcDNA3.1(+) vector expressing each of the four E proteins, SARS-CoV-2 N protein, SARS CoV-2 S protein, SARS CoV S protein, or HIV-1 Vpu and a vector expressing human BST-2 protein. At 30 h post-transfection, cells were starved for methionine/cysteine and radiolabeled with 500 μCi of ^35^S-methionine/cysteine for 16 h. Cell lysates were prepared in RIPA buffer and one third of the lysate was used to immunoprecipitate BST-2 proteins using an anti-BST-2 antibody; one third of the lysate was used to immunoprecipitate E proteins and Vpu (**Panel B**); S proteins (**Panel C**) using an anti-Spike antibody; or the N protein using an anti-GFP antibody (**Panel D**); and one third of the lysate was used to immunoprecipitate GAPDH to normalize loading as described in the Materials and Methods section.

We next examined the steady-state levels of BST-2 in the presence of the proteins analyzed above. HEK293 cells were co-transfected with vectors as described above and radiolabeled as described in the Materials and Methods section. Cell lysates were prepared, and the BST-2 proteins and viral proteins were immunoprecipitated with appropriate antibodies as described in the Materials and Methods section. The results indicate that in the presence of SARS-CoV and SARS-CoV-2 S proteins and HIV-1 Vpu, levels of immunoprecipitated BST-2 were significantly less than from cells transfected with the vector expressing BST-2 alone or cells co-transfected with vectors expressing BST-2 and the E proteins or SARS-CoV-2 N protein (**Fig. 9**). Taken together, the E and SARS-CoV-2 N proteins had no effect on BST-2 expression, while SARS-CoV and SARS-CoV-2 S proteins, and Vpu caused the downregulation of BST-2 from the cell surface and a decrease in total amounts of BST-2.

## DISCUSSION

The E protein of coronaviruses is a multi-functional protein that has several roles in the virus replication cycle. It is localized to the ER, ERGIC and Golgi compartments of the cell and associates with the M protein at the ERGIC during virus maturation (**32–35**). The palmitoylation of the conserved cysteine residues of the MHV E protein is important as substitution of these cysteine residues results in an unstable protein, aggregation of the M protein, and leads to incompetent virus-like particle (VLP) formation (**33, 36**). Other studies have shown that the E protein of infectious bronchitis virus (IBV) alters the secretory pathway by disrupting Golgi organization leading to the production of larger vesicles capable of transporting virions (**37–39**). The role of the E protein in virus maturation varies with the coronavirus. The deletion of the E gene of transmissible gastroenteritis virus (TGEV) completely arrests virus maturation while MHV, SARS-CoV and HCoV-OC43 expressing non-functional E proteins generate less infectious virus (**14, 40–42**). Replacement of the hydrophobic domain of IBV E with that of VSV G protein negatively affected virus maturation (**38**).

Here, we examined the biological properties of the SARS-CoV-2 E protein in the context of HIV-1. HIV-1 expresses the Vpu protein, which despite having a different sequence, has a similar length, an uncleaved leader sequence, and a similar membrane orientation as coronavirus E proteins. Additionally, both the E protein and Vpu enhance the release of their respective viruses. Several RNA viruses encode for proteins known as viroporins, which are virally encoded pore-forming membrane proteins that can alter membranes and ion homeostasis. These proteins are important in entry and/or egress from cells (**43–46**). The prototypical viroporin is the M2 protein of influenza A virus and since the identification of its ion channel properties, viroporins have been identified in several different viruses including HIV-1 (Vpu), hepatitis C virus (p7 protein), respiratory syncytial virus (SH protein), classical swine fever virus (p7 protein), and human papillomavirus (E5 protein) (**47–54**). The E proteins from SARS-CoV, MERS-CoV, and IBV have been reported to be viroporins while there are no reports of an ion channel activity associated with the HCoV-OC43 E protein (**25, 55–61**). Viroporins have been shown to transport different ions but generally are weakly selective for cations (H^+^, K^+^, Na^+^, Ca^++^) over anions and can alter host cell homeostasis at various stages during infection, often involving viral entry and/or release (**62**). However, other cellular functions such as apoptosis and vesicular trafficking can also be perturbed by viroporins (**63**). In a recent study, investigators showed that the M protein and E proteins of SARS-CoV-2 regulated the intracellular trafficking of the S protein at the site of assembly (**64**). These investigators showed that the M protein interaction with the C-terminal domain of the S protein mediated retention of the S protein in the ERGIC whereas the E protein caused retention of the S protein by modulating the cell secretory pathway (**64**). Our results revealed the SARS-CoV-2 (and SARS-CoV) E proteins significantly inhibited the unmodified HIV-1 and HIV-1Δ*vpu* infection while the E proteins from MERS and HCoV-OC43 were less restrictive.

Mechanistically, we showed that neither viral integration nor RNA synthesis were impaired by the presence of SARS-CoV-2 E protein. We observed that in the presence of the SARS-CoV-2 E protein, the levels of viral protein synthesis were significantly reduced. Previously, the SARS-CoV E protein was shown to be a channel for calcium ions via its viroporin activity (**65**). The ER is the major store of Ca^++^ within eukaryotic cells and is involved in lipid biosynthesis, detoxification of chemicals, and *de novo* protein synthesis and protein folding. Within the ER, molecular chaperones such as Bip/GRP78 are tasked to properly fold proteins such that they can transit to the Golgi complex. Thus, the ER provides a special environment for these chaperones for proper protein folding. Under conditions of Ca^++^ depletion, the ability of calcium-dependent molecular chaperones to properly fold proteins is reduced, which leads to the accumulation of misfolded proteins, ER stress, and initiation of an unfolded protein response (UPR). The UPR is a cellular pathway that detects and attempts to alleviate ER stress (**66, 67**). The UPR is initiated by the activation of three canonical ER stress sensors; the PKR-like endoplasmic reticulum kinase (PERK), activating transcription factor 6 (ATF6), and inositol-requiring enzyme 1α (IRE1-α) (**66–71**). Here we show that expression of the SARS-CoV-2 E protein caused the phosphorylation of eIF-2α to attenuate general protein synthesis. Phosphorylation of eIF-2α allows the translation of ATF4 mRNA, which encodes a transcription factor controlling the transcription of genes involved in autophagy, apoptosis, amino acid metabolism and antioxidant responses. In a previous study that examined gene expression in VeroE6 and MA104 cells infected with SARS-CoV and SARS-CoVΔE, investigators showed that SARS-CoV E protein down-regulated the inositol requiring enzyme (IRE-1) signaling pathway of the unfolded protein response (UPR) pathway but not the PERK or ATF6 pathways (**72**). They also found that expression of the E protein in *trans* reduced the stress response in cells infected with SARS-CoVΔE (**72**). There could be several reasons for the differences in the results of the two studies. These include the: a) cell types analyzed; b) transfection of cells with vector expressing a single protein from SARS-CoV-2 versus the use of wild type SARS-CoV and SARS-CoV E-deleted viruses; c) differences in the genome structure of SARS-CoV-2 and SARS-CoV; and d) differences in the amino acid sequences of the two E proteins (4 amino acid substitutions). Thus, this is the first report of the SARS-CoV-2 E protein involved in ER stress and phosphorylation of eIF-2α (**see accompanying paper**). Our analysis suggested that the decreased HIV-1 protein synthesis in cells expressing the E protein was correlated with the phosphorylation of eIF-2α. We show that indeed a decrease in protein synthesis occurs in E protein transfected cells (**see accompanying paper**).

Another possible mechanism for the observed decrease in HIV-1 protein expression may be related to autophagy. Autophagy, first discovered in yeast, is a multistep degradative process that maintains cell homeostasis by eliminating misfolded/old elements (proteins and organelles) trapped in autophagosomes targeted to fuse with lysosomes to obtain nutrients (**73,74**). Autophagy is also relevant in the innate and adaptive immunity against viral infections (**75–78**). HIV-1 infection has been shown to induce (macrophages) or inhibit (CD4+ T cells) autophagy (**78–79**). During the initial entry step, the HIV-1 gp120/gp41 proteins at the surface of the virus bind to CD4 receptors and co-receptor CCR5 or CXCR4, initiating autophagic response in CD4+ T cells. This autophagic process represents an anti-HIV-1 response by the host cell leading to the selective degradation of HIV-1 Tat (**80–82**). In later stages of HIV-1 infection there is an increased activation of mechanistic target of rapamycin complex 1 (mTORC1) (a regulator of autophagy) which leads to an inhibition of autophagy (**83**). Additionally, HIV-1 Nef protein interacts with Beclin 1 and its inhibitor BCL2 (**84**). In our accompanying paper, we show that the expression of the E protein leads to increased lipidation of microtubule-associated protein 1A/1B-light chain 3 (LC3) to yield LC3-II, which is required for autophagy. This could lead to the degradation of the viral proteins and explain our observations.

We also examined the E proteins from SARS-CoV-2, SARS-CoV, MERS-CoV, and HCoV-OC43 for the ability to down-regulate bone marrow stromal antigen-2 (BST-2; also known as CD317 or tetherin). This type II transmembrane protein was first discovered as a cell surface marker for differentiated and neoplastic B cell types (**85**) and later was shown to be induced by type I and II interferons. It is now understood that BST-2 activates NF-κB resulting in the activation of IFN-I responses and proinflammatory responses against viruses (**86**). The antiviral functions of BST-2 were first demonstrated with HIV-1 that lacked the *vpu* gene (**20, 87**). Since its initial identification as an antiviral restriction factor, BST-2 has been shown to tether several enveloped viruses including coronaviruses. BST-2 was shown to inhibit the release of HCoV-229E although the protein responsible for overcoming this restriction factor was not investigated (**88**). Later, it was shown that the SARS-CoV S protein caused the degradation of BST-2 (**89**). We showed that unlike Vpu, the E proteins neither down-regulated BST-2 expression at the cell surface nor caused BST-2 degradation while the S protein of SARS-CoV and SARS-CoV-2 clearly down-regulated surface expression of BST-2 and was accompanied by a decrease in total amounts of BST-2.

In conclusion, we showed that the E protein viroporin from SARS-CoV-2 and SARS-CoV potently restricted HIV-1 infection. Our data show that the SARS-CoV-2 E protein activated an unfolded protein response leading to the phosphorylation of eIF-2α, which is known to attenuate protein synthesis. These results show for the first time that a viroporin from one virus can inhibit the infection by another virus. It will also be of interest to determine what domain(s) of the S protein is responsible for the S/BST-2 interactions.

## MATERIALS AND METHODS

### Cells, viruses, and plasmids

HEK293 cells were used for transfection of vectors expressing coronavirus proteins and the HIV-1 genome. The TZM-bl cell line was used as an indicator to measure HIV-1 infectivity (**90–95**). Both cell lines were maintained as previously described (**29, 30, 90**). Plasmids with the entire HIV-1 NL4-3 genome (pNL4-3) and pNL4-3Δ*vpu* were obtained from the NIH AIDS Reagent Branch. Plasmids (all pcDNA3.1(+) based) expressing E proteins (SARS-CoV-2 E: accession #QIH45055, SARS-CoV E: accession # AAP13443, MERS-CoV E: accession #ATG84849, and HCoV-OC43 E: accession #ARA15423), and the HIV-1 Vpu protein (strain NL4-3) were synthesized by Synbio Technologies with C-terminal HA-tags and were sequenced to ensure that no deletions or other mutations were introduced during the synthesis. Plasmids expressing the SARS-CoV and SARS-CoV-2 S proteins were purchased from Sino Biologicals (catalog # VG40150-G-N; VG40589-CY). Expression of different coronavirus E proteins, the SARS-CoV-2 S, and SARS-CoV S protein were confirmed by transfection with the Turbofect transfection reagent (ThermoFisher) followed by radiolabeling and immunoprecipitation analysis using a mouse monoclonal antibody directed against the HA-tag (Thermo-Fisher, #26183) or an antibody against SARS-CoV Spike protein (Sino Biologicals, #40590-T62).

### Immunofluorescence studies

To examine the intracellular localization of the SARS-CoV-2 E protein, COS-7 cells grown on 13 mm cover slips were transfected with either the empty pcDNA3.1(+) vector or one expressing the SARS-CoV-2 E protein using Turbofect transfection reagent (ThermoFisher). At 24 h pt, cells were washed in PBS, fixed in 4% paraformaldehyde in PBS for 15 min, permeabilized with 1% Triton X-100 in PBS, and blocked for one hour with 22.5 mg/mL glycine and 1% BSA in PBST. The cultures were then incubated at 4C overnight with an and a rabbit monoclonal antibody against either Golgin-97 (Abcam, #ab84340) or Calnexin (Cell Signaling Technology, #2679). The cells were washed in PBS, incubated with a secondary goat anti-rabbit antibody conjugated to AlexaFluor™-488 (Invitrogen, A11008) for 1 h at room temperature, washed, and reacted with rabbit anti-mouse conjugated to AlexaFluor™- 594 (Invitrogen, A11062) for 1 h. Cells were counterstained with DAPI, and the coverslips were mounted on glass slides with a glycerol-containing mounting medium (SlowFade antifade solution A; Invitrogen). Coverslips were viewed with a Leica TCS SP8 Confocal Microscope with a 100X objective and a 2X digital zoom using the Leica Application Suite X (LASX) as previously described. A minimum of 100 cells were examined for each sample, and the results presented in the figures are representative of each sample.

### Analysis of infectious HIV-1 production in the presence of E proteins

To analyze the virus restriction properties of the E proteins, HEK293 cells were transfected with either the empty pcDNA3.1(+) vector, a vector expressing gD (a positive control for restriction), gD[ΔTMCT] (a negative control for restriction) or E proteins and pNL4-3 (**29, 30, 90**). At 48 h post-transfection (pt), the culture medium was collected, clarified by low-speed centrifugation, and the supernatants analyzed for infectious virus by titration on TZM-bl cells (**29, 30, 91–95**). All assays were performed at least four times and analyzed for statistical significance using two-tiered Student’s *t*-test with cells co-transfected with the empty pcDNA3.1(+) and pNL4-3 set at 100% infectivity.

### Analysis of infectious HSV-1 produced in the presence of SARS-CoV-2 E protein

To determine if the SARS-CoV-2 E protein would restrict the replication of HSV-1, HEK293 cells were transfected with either the empty vector or with a vector expressing SARS-CoV-2 E protein. At 48 h, cells were infected with HSV-1 (0.01 pfu/cell) for 2 h. The cells were collected at 24, and 48 h post-infection, and virus progeny production was determined by titration on Vero cells. Briefly, sterile skim milk was added to the culture, the cells were scraped into the medium, and briefly sonicated. Levels of infectious virus were determined by preparing a series of 10-fold dilutions of the culture supernatant followed by inoculation of Vero cells. The number of plaque forming units were determined by standard procedures.

### Biosynthesis and processing of viral proteins in the presence of E proteins

The biosynthesis and processing of HIV-1 proteins were examined in the presence of the coronavirus E proteins. HEK293 cells were co-transfected with empty pcDNA3.1(+) or the vector expressing E proteins and pNL4-3. At 30 h, the cells were washed and incubated in DMEM without methionine/cysteine for 2 h. The cells were washed and radiolabeled in DMEM containing 500 μCi ^35^S-Translabel (methionine and cysteine, Perkin-Elmer) for 16 h. Cell lysates were prepared, and culture medium processed as previously described (**29, 30, 90**). HIV-1 proteins were immunoprecipitated using a cocktail of antibodies previously described and is referred to in the figures as “anti-HIV” antibodies (**29, 30, 90**). The E proteins were immunoprecipitated using a monoclonal antibody directed against the HA-tag. Immunoprecipitates were collected by incubation with protein A-Sepharose beads overnight at 4C, the beads were washed with RIPA buffer, and the samples resuspended in sample reducing buffer. The samples were boiled, proteins separated by SDS-PAGE (10% or 12.5 % gels), and proteins visualized using standard radiographic techniques.

### Analysis of HIV-1 genome integration and viral RNA synthesis

To determine if the E proteins interfered with viral genome integration, we assessed the level of integration by amplification Alu repeat/*gag* sequences using the previously described procedure (**96**). HEK293 cells were transfected with either the empty pcDNA3.1(+) vector or one expressing SARS-CoV-2 E protein. At 24 h, equal levels of HIV-1ΔEnv-/VSV-G pseudotyped virus (M.O.I. of 0.1) were used to infect HEK293 cells for 48h. Total cellular DNA was purified using DNeasy Blood & Tissue Kit from Qiagen. With each experiment, a standard curve of the amplicon being measured was run in duplicate ranging from 10 to 1×10^11^ copies plus a no-template control. After initial incubations of 95C for 30 s, 40 cycles of amplification were carried out at 5 s at 95C, 34 s at 60C. Reactions were analyzed using the ABI Prism 7500 sequence detection system (PE-Applied Biosystems, Foster City, California).

For analysis of the HIV-1 transcription, HEK293 cells were transfected with either the empty pcDNA3.1(+) vector or one expressing the SARS-CoV-2 E. At 24 h post-transfection, cells were infected with pseudotyped HIV-1ΔE/VSV-G at MOI of 0.1. At 48 h post-transfection, cells were washed, pelleted and the RNA was extracted using Invitrogen PureLink RNA Extraction kit. The RNA was treated with PureLink DNase I for 15 min at room temperature followed by heat inactivation. The RNA was reverse transcribed using SuperScript IV RT kit and oligo(dT) 15 primer. The mixtures were treated with RNase H for 30 minutes at 37 C. Real-time quantitative PCR was performed using the procedures previously described primers: Tat forward, ES2440: 5’- GTCAGCCTAAAACTGCTTGTACCA-3’; Tat reverse, ES2445: 5’-GCCTGTCGGGTCCCCTC-3’; and Tat probe, MH603: 5’-(FAM)-CTCCTATGGCAGGAAGA-(TAMRA)-3’. A standard curve of the RNA amplicon was run ranging from 10 to 1×10^11^ copies. Reactions containing 1x premix (Takara), 0.5μM forward primer, 0.5μM reverse primer, 0.25μM probe and 1x RoxDyeII in a 20μl volume. The thermal cycle is 95 C for 30 s, followed by 40 cycles of amplification for 5 s at 95C and 34 s at 60C. Reactions were analyzed using the ABI Prism 7500 sequence detection system (PE-Applied Biosystems, Foster City, California).

### Analysis of the phosphorylation of eIF-2α

To determine if the expression of the E protein of SARS-CoV-2 resulted in phosphorylation of eIF2α, HEK293 cells seeded in 6-well plates were either left untransfected or transfected with 1 μg of pUC19 or of a SARS-CoV-2 E-expressing plasmid. The cells were harvested at 48 h pt and equal amounts of proteins were analyzed by western blot for p-eIF2α or total eIF2α (both antibodies were obtained from Cell Signaling). Expression of E was detected using an HA-tag antibody (Invitrogen). β-actin was used as a loading control.

### Analysis of BST-2 down-regulation by various E proteins and SARS-CoV-2 S, and N proteins

We determined if the E proteins from several coronaviruses (SARS-CoV-2, SARS-CoV, MERS-CoV, HCoV-OC43), the SARS-CoV S protein, and SARS-CoV-2 S, M, and N proteins were capable of down-regulating cell surface bone marrow stromal antigen 2 (BST-2). HEK293 cells were co-transfected with either empty pcDNA3.1(+) vector and pcDNA3.1(+) expressing the human BST-2 protein, or pcDNA3.1(+) expressing each of the proteins described above and BST-2. Cells co-transfected with vectors expressing HIV-1 Vpu and BST-2 was used as a control for BST-2 down-regulation. At 48 h, the cells were stained with a mouse monoclonal antibody directed against BST-2 and subjected to flow cytometric analysis using a BD LSR II flow cytometer. Aliquots of cells from the same co-transfections were also analyzed for protein expression by immunoblots using antibodies directed against the HA-tag (E proteins, SARS-CoV-2 S protein, and Vpu), the SARS-Cov-1 S protein, or against GFP to monitor SARS-CoV-2 N-GFP expression.

To analyze if the above proteins led to a steady state reduction in BST-2, HEK293 cells were co-transfected with vectors expressing either SARS-CoV and SARS-CoV-2 S proteins, N protein tagged with EGFP (at the C-terminus), the four E-HA proteins, or HIV-1 Vpu-HA and a vector expressing BST-2. At 36 h post-transfection, cells were starved of methionine/cysteine and radiolabeled for 2 h in methionine/cysteine-free media containing 500 μCi of ^35^S-methionine/cysteine for 16 h. Cells were washed, lysed in 1X RIPA buffer and clarified by centrifugation. The lysates were transferred to new tubes and BST-2 (using an anti-BST-2 antibody), the E proteins (all having a C-terminal HA-tag), and SARS-CoV and SARS-CoV-2 S proteins, and N-GFP proteins, or GAPDH (to equalize loading) immunoprecipitated using appropriate antibodies. The immunoprecipitated proteins were washed in RIPA buffer and boiled in sample reducing buffer. The proteins were separated using SDS-PAGE and visualized by standard radiographic techniques.

## ACKNOWLEDGMENTS

This research was funded by the NIH/NIAID 1R21AI158229-01 to E.B.S and M.K. and the KUMC Frontiers Clinical and Translational Science Awards Program, grant number 5UL1TR002366-04. We thank the ACGT for the sequence analyses and the KUMC Biotechnology Core for oligonucleotide synthesis. The following reagents were obtained from the NIH AIDS Reagents Branch, Division of AIDS, NIAID, NIH: 1) the HIV-1 p24 Gag monoclonal (catalog #ARP-6521) from Dr. Michael H. Malim; 2) pNL4-3 (catalog #ARP-114) was obtained from Dr. Malcolm Martin and Klaus Strebel; 3) the HIV-1 BaL.26 clone (Cat# ARP-11446) from Dr. J.R. Mascola; 4) the goat anti-gp160 antibodies (Catalog # ARP-51, ARP-189, ARP-191).

